# Streptococcal superantigen SpeC induces IL-8 secretion in human epithelial cells

**DOI:** 10.64898/2026.05.18.725648

**Authors:** Rige Na, Siyu Guo, Xiaolan Zhang

## Abstract

Streptococcal pyrogenic exotoxin C (SpeC) is a prototypical superantigen produced by group A *Streptococcus*. It potently activates a broad subset of T lymphocytes via a bridging interaction involving TCRβ−SpeC−MHC-II. Our recent work demonstrated that SpeC induced profound release of IL-8 from human pharyngeal epithelial cells and this effect was reversible through a specific point mutation in SpeC. This study systematically investigated cellular signaling pathways using integrated transcriptomic profiling and Western blot analysis, with a focus on membrane-associated receptors and downstream intracellular signaling effectors. Our results demonstrate that this biological process is critically associated with the activation of Erk1/2, p38 MAPK and NF-κB signaling cascade. This study identifies a novel mechanism through which a bacterial superantigen target epithelial cells−the body primary physical barrier and first line of innate immune defense.

## 1 Introduction

Group A Streptococcus (GAS) is the most clinically significant and virulent streptococcal pathogen in humans, capable of causing a broad spectrum of diseases ranging from superficial mucocutaneous infections (e.g., pharyngitis and scarlet fever) and localized skin infections (e.g., impetigo) to severe invasive conditions such as necrotizing fasciitis and streptococcal toxic shock syndrome. Additionally, GAS infection may trigger post-infectious autoimmune sequelae, most notably acute rheumatic fever and its long-term complication, rheumatic heart disease ^[1]^. Superantigens (SAgs) are a class of potent T-cell mitogens predominantly secreted by bacterial pathogens. Unlike conventional antigens, SAgs bypass normal antigen processing and presentation pathways, leading to the polyclonal activation of a substantial proportion of T lymphocytes and thereby triggering excessive cytokine release and immune dysregulation associated with severe inflammatory pathologies. GAS produces at least 11 structurally and functionally distinct SAgs, including streptococcal pyrogenic exotoxins A and C (SpeA, SpeC), SpeG through SpeM, streptococcal superantigen (SSA), and streptococcal mitogenic exotoxin Z (SmeZ). The capacity of these SAgs to subvert host adaptive immunity represents a central mechanism underlying GAS virulence and pathogenesis ^[2]^. Epidemiological studies have demonstrated that the carriage rate of the *spec* gene among different isolates recovered from pharyngeal and tonsillar specimens in multiple regions of China exceeds 90%. SpeC is a non-glycosylated, low-molecular-weight extracellular protein. It exhibits remarkable stability under extreme conditions, including high temperature, low pH, desiccation, and proteolytic degradation, and is capable of inducing pyrogenic responses in humans ^[2]^. The *spec* gene encoded by MGAS9429, comprises 708 base pairs and is located within a prophage element, facilitating horizontal transfer among GAS strains. Its nucleotide sequence is identical to those of *spec* alleles in other well-characterized GAS strains including SF370, MGAS8232, and MGAS6180, yet exhibits only 24% nucleotide sequence identity with *speA* ^[3, 4]^.

SpeC functions as a superantigen that rapidly activates a large population of T cells. Its mechanism involves bridging the T cell receptor (TCR) β chain on αβ T lymphocytes with the outer domain of major histocompatibility complex class II (MHC-II) molecules expressed on antigen-presenting cells—forming a ternary TCR β-SpeC-MHC-II complex. SpeC binds the α chain of MHC-II with low affinity, while exhibiting Zn^2+^-dependent high-affinity binding to the β chain of MHC-II ^[5]^. SpeC avoids direct interaction with the highly variable complementarity determining region 3 (CDR3) core region of the TCR β chain and instead preferentially engages the comparatively conserved CDR1 and CDR2 regions of the same chain. Consequently, its activity is independent of antigen specificity, enabling broad activation of diverse T cell populations with substantially enhanced efficiency—up to several orders of magnitude higher than antigen-dependent mechanisms ^[2, 5-7]^. This atypical and non-antigen-specific activation triggers sustained cytokine production by T lymphocytes and antigen-presenting cells, including IL-1β, IL-2, IL-6, IL-8, IL-10, IFN-γ, TNF-α, G-CSF, and IL-17, thereby inducing dysregulated immune responses. These characteristics are strongly associated with the pathogenic potential of SpeC in GAS-mediated scarlet fever and toxic shock syndrome ^[8]^.

Prior studies have established that SpeC activates T cells and elicits robust immune responses; however, its functional impact on non-T cell populations remains poorly characterized. Our findings demonstrate that SpeC induces IL-8 secretion in epithelial cells and modulates associated signaling pathways, suggesting a potential novel pathogenic mechanism underlying bacterial superantigen activity.

## 2 Materials and Methods

### 2.1 Bacterial Strains and Culture Condition

Group A *Streptococcus* (GAS) was cultured in Todd-Hewitt broth (BD Biosciences) supplemented with 0.2% (w/v) yeast extract (THY medium) at 37 ℃ under a humidified 5% CO_2_ atmosphere. Bacterial cultures were grown to mid-logarithmic phase (OD_600_ = 0.6-0.8) in THY, then harvested by centrifugation, washed with phosphate-buffered saline (PBS), and resuspended in PBS to the desired concentration.

### 2.2 Gene Synthesis, Protein Expression and Purification

The full-length SpeC protein sequence was chemically synthesized and cloned into the pET-30a(+) vector between the NdeI and HindIII restriction sites to generate the corresponding recombinant plasmid. Subsequently, the recombinant plasmid was transformed into *E*.*coli* BL21(DE3) competent cells. Protein expression was induced by adding isopropyl-β-D-thiogalactoside (IPTG), followed by incubation at 37 ℃ with shaking at 200 rpm for 4 h. Bacterial cells were harvested by centrifugation, resuspended in lysis buffer, and lysed by sonication. The cell lysate was then clarified by centrifugation and filtration. Recombinant SpeC protein present in the supernatant was purified using Ni^2+^-affinity chromatography. Protein purity and molecular weight were assessed by 12% SDS-PAGE, and protein concentration was quantified using a BCA protein assay kit.

### 2.3 Mouse Vaccination Experiments

Female BALB/c mice (6 to 8 weeks old) were purchased from Liaoning Changsheng Biotechnology Co., Ltd. The mice received intramuscular immunization three times with protein plus adjuvant or PBS plus adjuvant on days 0, 14 and 28. Body weight was measured every two weeks, and the sera were collected at specific time points. On day 42, mice were sacrificed by the cervical dislocation after being anesthetized with tribromoethanol (4 mg/10 g body weight) by intraperitoneal injection. All animal procedures were reviewed and approved by the Institutional Research Board of Harbin Medical University (Assurance Number: HMUIRB2024011).

### 2.4 Cytokine Quantification

The samples or homogenate underwent centrifugation to obtain supernatant, from which the concentrations of IL-8 were quantified using ELISA kits in accordance with the guidelines provided by the manufacturer (Elabscience Biotechnology Co., Ltd.).

### 2.5 Statistical Analysis

The data were analyzed using Student’s *t*-test or paired *t*-test. All figures were plotted using GraphPad Prism version 8. *P* values less than 0.05 were considered as statistically significant.

## 3 Results

### 3.1 Expression and purification of the SpeC recombinant protein and seven alanine-substituted mutants in a prokaryotic system

In this study, the SpeC sequence derived from MGAS9429 was selected as the template for structural modeling. Using the AlphaFold3 algorithm, we performed a high-confidence tertiary structure prediction of SpeC. Subsequently, site-directed mutagenesis was designed to introduce specific amino acid substitutions at seven evolutionarily and functionally conserved residues. Soluble SpeC expression was observed and was therefore selected as the standard induction condition for large-scale protein production (Figure 1A). The recombinant protein was purified via Ni^2+^-affinity chromatography, and the target protein was recovered in high purity through stepwise elution (Figure 1B). Using this approach, all seven SpeC alanine-substitution mutants— Y15A, F31A, Y76A, I77A, Y85A, Y87A, and R181A—were successfully expressed and purified to homogeneity. Western blot analysis demonstrated that the anti-His tag antibody specifically recognized a band at approximately 25 kDa, consistent with the predicted molecular weight of the recombinant protein (Figure 1C). In contrast, immunoblotting with SpeC-specific antiserum detected both wild-type and mutant proteins; however, the signal intensity differed significantly between the two variants. This differential reactivity suggests that the introduced point mutation may impair epitope integrity or alter antigenic conformation, thereby modulating immune recognition by the antiserum (Figure 1D).

**Figure 1.**
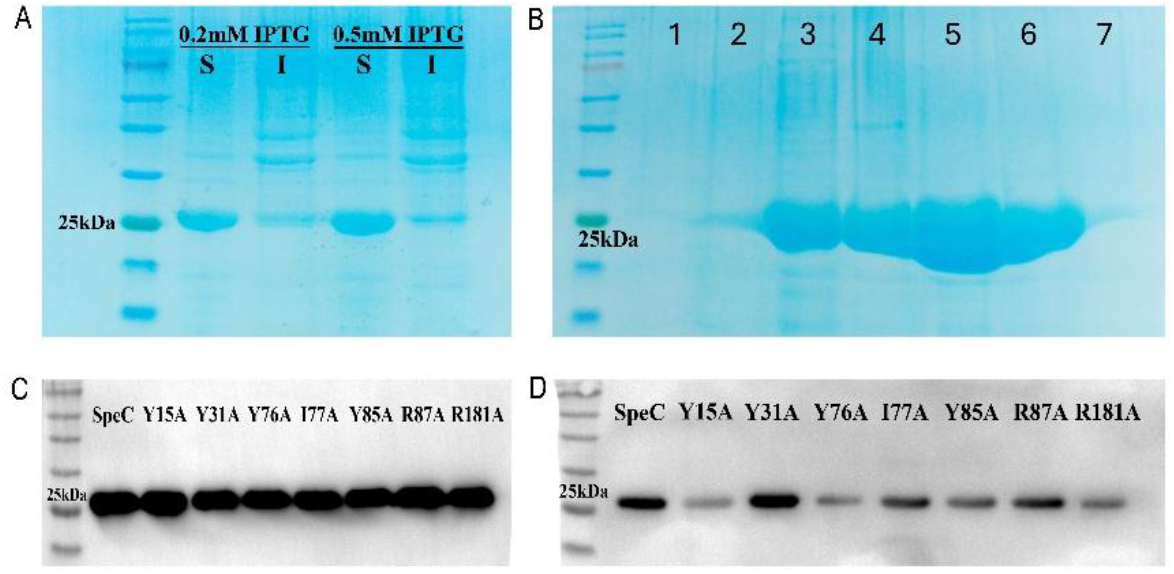
Qualitative analysis of the recombinant SpeC protein and seven site-directed mutants. (A) Coomassie brilliant blue staining confirmed the expression of the recombinant SpeC protein. S indicated supernatant and I indicated inclusion body. (B) Analysis of nickel-affinity chromatography purification. (C) Western blot analysis using anti-His tag antibodies. (D) Western blot analysis using SpeC-specific antiserum. © 2026 [Xiaolan Zhang]. Unauthorized reproduction or use is prohibited.

### 3.2 SpeC stimulation significantly enhances IL-8 secretion from pharyngeal epithelial cells, whereas all three site-directed mutants effectively abrogate this induction

Cells were stimulated with either recombinant SpeC protein or an equimolar amount of the corresponding point-mutated variant. The cytokine concentrations in the culture supernatants were quantified by enzyme-linked immunosorbent assay (ELISA). As shown in Figure 2, wild-type SpeC potently induced IL-8 secretion from cells. Notably, the Y15A, F31A, I77A, and Y85A point mutants also stimulated IL-8 release, albeit to varying extents relative to the wild-type protein. In contrast, the Y76A, Y87A, and R181A mutants failed to significantly enhance IL-8 secretion from the cells, indicating that these three residues are critical for the functional activity of SpeC. IL-8, a key neutrophil chemoattractant, plays a pivotal role in recruiting both innate and adaptive immune cells to sites of infection and contributes critically to the host immune defense response. The SpeC protein has been shown to induce IL-8 secretion in pharyngeal epithelial cells, indicating its potential involvement in immune evasion and/or host tissue pathology through modulation of host inflammatory signaling pathways.

**Figure 2.**
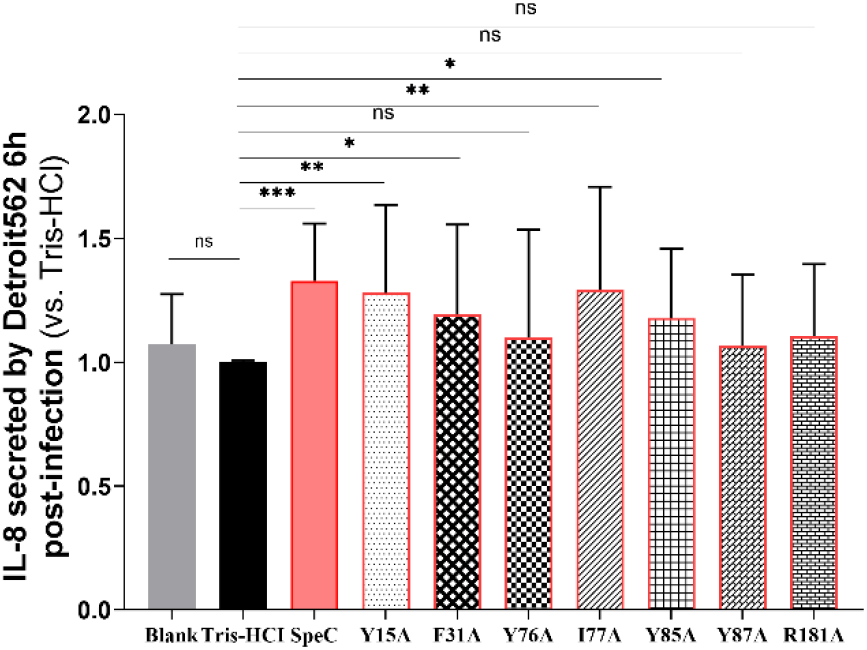
Normalized quantitative analysis of IL-8 concentrations in the culture supernatants of epithelial cells. IL-8 concentrations across experimental groups were normalized to the Tris-HCl control group, and results are expressed as mean ± SD, n=6, \**P*< 0.05, \*\**P*< 0.01, \*\*\**P*< 0.001. © 2026 [Xiaolan Zhang]. Unauthorized reproduction or use is prohibited.

### 3.3 SpeC can induce the phosphorylation of key signaling molecules—including Erk1/2, p38 MAPK, and NF-κB in pharyngeal epithelial cells

To investigate the molecular mechanisms underlying SpeC-induced IL-8 secretion in human epithelial cells, this study performed whole-transcriptome RNA sequencing on cells stimulated with SpeC relative to those treated with Tris-HCl buffer as a vehicle control. As illustrated in Figures 3A and 3B, SpeC stimulation induced differential expression of 78 upregulated and 19 downregulated genes relative to the control group. Gene set enrichment analysis revealed significant enrichment of these differentially expressed genes in multiple inflammation-associated signaling pathways, including the TNF, IL-17, and NF-κB signaling pathways. To validate the sequencing findings, Western blot analysis was performed to quantify the phosphorylation levels of key signaling molecules (Figure 3C-J). The results demonstrate that SpeC significantly enhances Erk1/2 phosphorylation, an effect completely abrogated by the Y87A and R181A mutations. SpeC also robustly induces p38 phosphorylation, which is abolished by the I77A, Y85A, Y87A, and R181A mutations. Moreover, SpeC promotes NF-κB phosphorylation, and this effect is likewise eliminated by the Y87A and R181A mutations. The aforementioned findings suggest that SpeC modulates the inflammatory response in epithelial cells through activation of the Erk1/2, p38 MAPK, and NF-κB signaling pathways; moreover, amino acid residues at positions 87 and 181 appear to be critical for the biological activities of SpeC.

**Figure 3.**
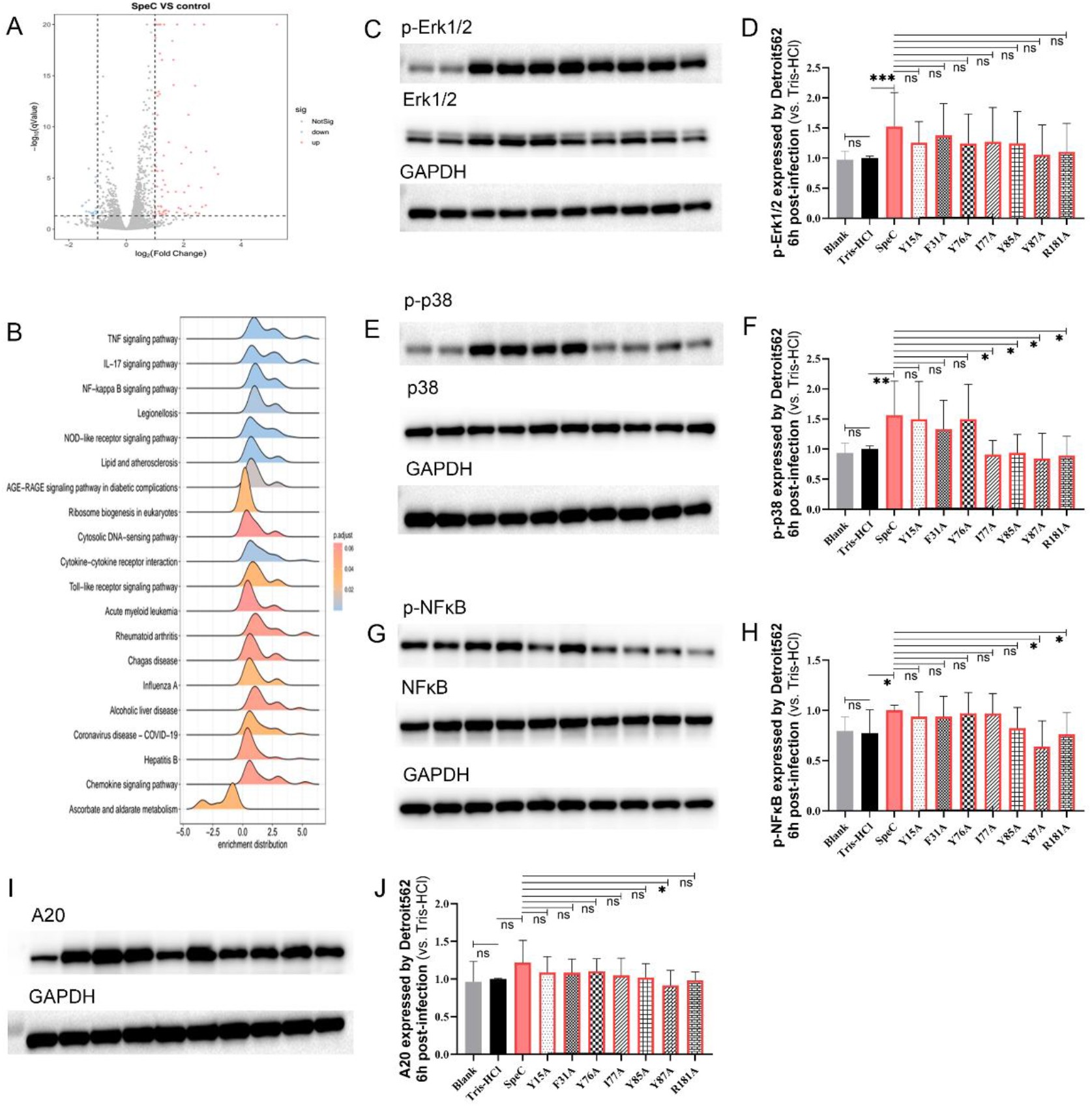
Transcriptome sequencing analysis and phosphorylation levels of signaling pathway molecules. (A) Volcano plot illustrating differentially expressed genes: up-regulated genes are highlighted in red, down-regulated genes in blue, and non-differentially expressed genes in gray. (B) Ridge plot of gene set enrichment analysis. The x-axis displays the log_2_-transformed expression values of core enriched genes across significantly enriched pathways; the y-axis represents the density distribution of enriched genes within each pathway. Significance is inversely proportional to the displayed value: lower (blue-shaded) values indicate higher statistical significance. (C, E, G, I) Western blot analysis was performed to assess the expression levels of Erk1/2, p38 MAPK, NF-κB, and A20. Sample lanes, from left to right, correspond to the following conditions: blank control, Tris-HCl buffer control, wild-type SpeC, and the point mutants Y15A, F31A, Y76A, I77A, Y85A, Y87A, and R181A. (D, F, H, J) Densitometric measurement of Western blot bands. © 2026 [Xiaolan Zhang]. Unauthorized reproduction or use is prohibited.

### Limitations

This preprint presents preliminary experimental findings. AI-assisted deep analysis of sequencing data is ongoing. Additional validation experiments are in progress to consolidate the proposed mechanism. The complete dataset and refined quantitative measurements will be included in the final manuscript.

## Data Availability

Raw transcriptome sequencing data and raw pictures will be deposited in GEO upon formal publication. Processed data, materials and pictures supporting this preprint are reserved and no part of this preprint may be reproduced. © 2026 [Xiaolan Zhang]. Unauthorized reproduction or use is prohibited.

## Ethics Statement

All animal studies were performed in strict accordance with the guidelines set by the university and all animal procedures were reviewed and approved by the Institutional Research Board of Harbin Medical University (Assurance Number: HMUIRB2024011).

## Author Contributions

Conceptualization, XZ; Methodology, XZ; Software, RN; Validation, SG; Formal analysis, XZ; Investigation, RN and SG; Writing, XZ; Supervision, XZ; Project administration, XZ; funding acquisition, XZ. All authors have read and agreed to the published preprint.

